# The Leiden *ex vivo* human growth plate model in severe tall stature: a proof-of-concept study

**DOI:** 10.64898/2026.08.25.746685

**Authors:** Margo Tuerlings, Yolande F. M. Ramos, Helena E. D. Suchiman, Sana S. Sayedipour, Sjoerd D. Joustra, Anika Rabelink-Hoogenstraaten, Hermine A. van Duyvenvoorde, Dagmar R. J. Kempink, Pieter Bas de Witte, Ingrid Meulenbelt, Christiaan de Bruin

**Affiliations:** Dept. of Biomedical Data Sciences, Section Molecular Epidemiology, Leiden University Medical Centre, Leiden, The Netherlands; Dept. of Paediatric Endocrinology, Willem-Alexander Children’s Hospital, Leiden University Medical Centre, the Netherlands; Dept. of Pediatrics, Spaarne Gasthuis, Haarlem, the Netherlands; Dept. of Orthopedics, Leiden University Medical Centre, Leiden, The Netherlands; Dept. Clinical Genetics, Leiden University Medical Centre, Leiden, The Netherlands; Dept. of Orthopedics & Sports Medicine, Erasmus Medical Centre / Sophia Children’s Hospital, Rotterdam, The Netherlands

**Keywords:** Human growth plate, Percutaneous epiphysiodesis, Chondrocytes, Endochondral bone growth

## Abstract

**Background:** Viable human growth plate (GP) tissue is rarely available for translational research, limiting direct investigation of human longitudinal bone growth and pediatric growth disorders. In this proof-of-concept study, we aimed to determine feasibility to establish a clinically integrated *ex vivo* human growth plate model using tissue obtained during routine percutaneous epiphysiodesis (PE) procedures in adolescents treated for extreme tall stature or leg length difference due to trauma.

**Methods:** Growth plate tissue and cells were collected during PE and processed using protocols adapted from established methods of human osteoarthritic cartilage collection within the RAAK study. Collected tissues and three-dimensional cartilage pellets generated from isolated cells were assessed by histology. Proliferation of GP-derived chondrocytes was compared with osteoarthritis-derived articular chondrocytes.

**Results:** Across consecutive surgical procedures, viable GP tissue could be obtained reproducibly including the node of Ranvier, with only few samples failing to yield cells and no contamination. Isolated GP chondrocytes expanded successfully in two-dimensional culture and showed a stronger early proliferative response compared with RAAK-derived chondrocytes. In addition, GP-derived cells formed threedimensional organoids and histology confirmed cartilage-like matrix deposition supporting their capacity to generate neo-cartilage tissue *in vitro*.

**Conclusion:** This study demonstrates feasibility to obtain, culture, and functionally assess viable human growth plate tissue from routine PE surgery. As such, the Leiden *ex vivo* human growth plate model provides a unique platform to study local mechanisms of endochondral bone growth, link genetic determinants of height to functional growth plate biology, and support future therapeutic research in pediatric growth disorders.

## INTRODUCTION

Growth concerns, particularly short stature, are among the most common reason for referral to an academic pediatric endocrinologist (*1*). Multiple underlying causes of short stature are known, such as growth hormone (GH) deficiency, hypothyroidism, chronic illness, genetic defects, or lack of catch-up growth after low birth weight and/or length. When none of these apply, children are labelled as idiopathic short stature (ISS ll; (*2-4*)). In the absence of effective targeted therapies, for most ISS cases recombinant growth hormone (GH) therapy is being applied although its benefit was found to be inconsistent (*5, 6*). Henceforth, patients with ISS may experience severely reduced adult heights (height < -2SD) accompanied by a significant reduction in well-being and psychological concerns (*7, 8*). At the other end of the spectrum are adolescents who suffer from severe tall stature, defined as height > 3SD, or leg length difference. While short stature can be caused by a multitude of causes, the causes and occurrence of extreme tall stature is fairly limited (*7*). Some of these cases are diagnosed with pathological hypersecretion of hormones (e.g. androgens, estrogens, thyroid hormone) or monogenetic and syndromic conditions. Nonetheless, most children with extreme tall stature have a polygenic familial history, hence a normal growth plate physiology but appear at the tall end of the height spectrum (*9*). Human height is shown to be a quantitative complex trait with high heritability (70-90%; (*10*)). Genetic studies that aim to identify genes that determine the genetic basis of such complex traits, are proven to deliver powerful hints on the etio-pathophysiological factors at play.

To identify genetic variants that explain the heritability of height, comprehensive genome-wide association studies (GWAs), including 5.4 million individuals, resulted in the identification of 12,111 independent height-associated genetic variants (*11, 12*). Nonetheless, in the absence of human growth plate tissues, functional follow-up studies aiming to unravel biological mechanisms by which associated genes affect height, have primarily been performed in mouse growth plate chondrocytes (*11*). These studies showed that variations in height coincide with genes and epigenetic modifiers that regulate growth plate chondrocyte transitions and/or their sequential cellular phenotypes during endochondral ossification (*13-16*). Nonetheless, due to important differences in timing and closure of rodent growth plates, translation of the genetic findings to drug target discoveries for patients with growth concerns has made little progress (*17*). Therefore, it is generally acknowledged that advancing translation of biomedical results via functional (genomic) follow-up studies, to novel druggable targets for skeletal dysplasias and growth disorders, requires viable pediatric human growth plate tissues (*18*). However, since growth plate biopsies are not part of routine clinical diagnostics, this tissue is rarely available (*19, 20*). Moreover, joint replacement surgeries that could in theory provide an alternative source of *ex-vivo* growth plate tissue, are extremely rare in the pediatric age range and will not supply a substantial flow of growth plate tissue for translational research (*21*).

Over the past 20 years, elective bilateral percutaneous epiphysiodesis (PE) around the knees has increasingly been used in the Netherlands as an approach to reduce adult height when severe tall stature is expected (>205 cm in men and >190 cm in women; (*22*)). During PE, the growth plates of femur and/or tibia are percutaneously disrupted using a drill and/or curette. This focal physeal damage induces growth arrest at the treated site, thereby eliminating the remaining growth potential of the concerning physis. PE has shown clear efficacy, low rates of long-term complications and overall high rates of patient satisfaction (*23*). Importantly, PE is mainly performed in adolescent patients who are in the midst of their pubertal growth spurt, when growth plate chondrocytes are still highly active.

In this study, we set up a procedure to harvest intact primary human growth plate tissues during routine percutaneous PE procedures. Harvest tissues are then processed and stored following the previously established protocol of articular cartilage within the RAAK-study (*24*), hence eligible for functional followup studies including 3D *in-vitro* organoid (*25*), and *ex-vivo* tissue cultures (*26*), histology, and molecular epidemiological approaches (*27*). Consequently, the Leiden GROWTH study is designed to advance understanding of the biological processes at play in the growth plate. The current manuscript documents the design of the study, patient inclusion, the operating procedures, and the exploratory biomedical techniques that were used to assess the feasibility of this novel *ex vivo* growth plate model.

## METHODS

### Ethical approval and procedural safety

The study protocol was approved by the LUMC Medical Ethical Committee (P19.105), and all patients (and their parents if age was below 16 years) gave written informed consent. From the viewpoint of patient safety, it is important to note that during PE surgery the growth plates were biopsied with an 8 Gauge Jamshidi needle using standard PE incision, trajectory, and location. The caliber of the biopsy needle (8 Gauge) in this study was smaller or similar to generally used drills and curettes and the trajectory of the biopsy needle was the same as the trajectory used for the drill in a standard epiphysiodesis. Hence, the risk of post-operative complications as a result of this study was projected to be absent.

### Patient inclusion

Patients were enrolled at the pediatric orthopedic out-patient clinic at LUMC (Leiden). All patients were evaluated pre-operatively by a pediatric endocrinologist and a pediatric orthopedic surgeon as part of routine patient care. If a monogenetic cause for tall stature was suspected, patients received genetic evaluation using a targeted gene panel. Patients with a minimal predicted adult height >205 cm for males and >190 cm for females, whose growth plates were still sufficiently open for height reduction, were considered for PE using shared decision making. We also included children with normal stature that underwent PE for significant leg length difference. In these latter patients the orthopedic surgeon performed PE on the growth plate of the longer leg to reduce the leg difference. These growth plates served as internal controls.

Data were collected from patient records. Reference data were used to calculate body mass index (BMI; (*28*)), sitting height/height index (*29*), and target height (*30*). Bone age assessment was estimated in all patients for adult height prediction using standardized BoneXpert software V3.1 (*31*) and traditional Greulich & Pyle assessment by the pediatric endocrinologist (*32*). If clinically relevant, serial bone age assessments or additional X-rays of growth plates around the knees were obtained. Parental heights were obtained by history during the clinical encounters or measured when these heights were unknown.

### Surgical procedure

On the day of PE surgery, EDTA blood was obtained during perioperative routine IV placement for DNA extraction and storage for future genetic studies related to growth plate function. Surgery was performed under general anesthesia, by experienced pediatric orthopedic surgeons (DK and/or PBdW). The lateral and/or medial approach was applied with drilling for each distal femur and/or proximal tibia growth plate as commonly done. Pathology samples were acquired using an 8 Gauge Jamshidi needle. With the optimized approach, the Leiden method, this was done at the level of the node of Ranvier. Tissue biopsies and physeal drill debris were collected for both histology and isolation of cells. The biopsies were placed in normal saline before being transported to the research lab for collection within 2 hours. Upon arrival in the laboratory, growth plate tissue was dissected in separate layers to collect the different cell types. Isolated primary chondrocytes were cultured according to previously reported methods applied to the Research Articular osteoArthritis Cartilage (RAAK) study (*24, 33*) and used for the different experiments while checking validity and feasibility of our research model. This included cell proliferation and differentiation analyses, histology, measurement of various paracrine factors (such as CNP), and basic functional studies including (epi)genetic and transcriptomic (RT-qPCR) analyses to determine the effects of established growth-related peptides (such as GH and GH-antagonists) on human chondrocyte function *in vitro*.

### Postoperative cell and tissue processing procedure

The materials collected during PE surgery were processed within two hours post-surgery. If the biopsy sample was sufficiently large, it was divided into three parts and processed as follows: (1) one part was snapfrozen and stored in liquid nitrogen for future RNA/DNA isolation, (2) one part was fixed in 4% formaldehyde, decalcified in 10% EDTA, embedded in paraffin and stored for further analysis, and (3) one part was used to isolate chondrocytes as described previously.

in short, to isolate cells the biopsy was minced into small fragments and incubated overnight in expansion medium (DMEM (high glucose; Gibco), supplemented with 10% fetal bovine serum (FBS; Gibco), 100 U/ml penicillin and 100 ug/ml streptomycin (Gibco), 0.5 ng/ml FGF-2 (PeproTech)), in addition of 2 mg/mL collagenase type I. The physeal drill debris was incubated likewise but without further mincing. Next day, the cell suspension was resuspended and passed through a 40 μm mesh filter to remove any undigested tissue fragments. Cells were then reseeded in expansion medium. Upon reaching 80% confluency, the cells were collected, frozen (using 10% DMSO, Sigma), and cryopreserved for future use.

### 3D cartilage organoids

For 2D proliferation assay and generation of 3D organoids cells were passaged twice at approximately 80% confluency. For the proliferation assay, cells were harvested and counted (NucleoCounter NC-200, Chemometec) 24 and 96 hours after seeding. Remaining cells were collected in expansion medium by centrifugation (4 minutes, 1200 rpm; 2.5x10^5^ cells per organoid). After 24 hours, the expansion medium was replaced with chondrogenic differentiation medium (DMEM (high glucose; Gibco), supplemented with Ascorbic acid (50 μg/ml; Sigma-Aldrich), L-Proline (40 μg/ml; Sigma-Aldrich), Sodium Puryvate (100 μg/ml; Sigma-Aldrich), Dexamethasone (0.1 μM; Sigma-Aldrich), ITS+, 100 U/ml penicillin and 100 ug/ml streptomycin (Gibco) and TGF-β1 (10 ng/ml; PeproTech)), as described previously (*33*). Medium was refreshed every 3-4 days for 21 days. After this, the organoids were harvested and either fixed overnight with 4% formaldehyde and embedded in paraffin for histology, or lysed with Trizol for RNA isolation and gene expression analysis.

### Histochemistry

Embedded tissues, from either growth plate biopsies or 3D cartilage organoids, were sectioned (5 μm thickness). The sections were deparaffinized and rehydrated, then stained with either Alcian Blue to visualize glycosaminoglycans (GAGs) or with hematoxylin and eosin (H&E) to visualize the tissue structure.

### RNA isolation and gene expression analysis with RT-qPCR

RNA was isolated from the 3D cartilage organoids using the RNeasy Mini Kit (Qiagen) and cDNA was synthesized with the First Strand cDNA Synthesis Kit (Roche Applied Science), both following the manufacturer’s protocols. RT-qPCR was performed using SYBR Green without ROX reference dye (Roche Applied Science) and measurements were done using QuantStudio 6 Real-Time PCR (Applied Biosystems). Gene expression levels were corrected for the housekeeping genes *GAPDH* and *SDHA* and reported as -ΔCT values.

## RESULTS

### Patient characteristics

To date, a total number of 28 patients were enrolled in our study. Baseline patient characteristics are presented in **Supplementary Table S1**. In total 15 out of 28 patients had a diagnosis of idiopathic, nonsyndromic familial tall stature and seven patients presented with normal stature, but underwent unilateral PE due to leg length difference. Six patients had genetically confirmed, syndromic tall stature (5 cases of Marfan syndrome, 1 patient with Weaver syndrome). At the time of PE the mean bone age was 13.8 years among males and 12.5 years among females. This age coincides with the known peak of growth spurt in most adolescent and could imply that the chondrocytes were highly active at the time of harvesting.

### Percutaneous epiphysiodesis and histology of growth plate tissue

We first sought to optimally collect intact growth plate tissue during the PE surgery by comparing different biopsy approaches aiming to obtain Ranvier’s node and/or a transverse architectural layers of GP using a Jamshidi needle (**Supplementary Figure S1**). As compared to the standard collection method taken with the Jamshidi needle, visualization of intact growth plate tissues including Node of Ranvier was improved upon applying the Leiden method. Histological analysis of the latter with Haematoxylin Eosin (HE) and Alcian Blue staining featured different growth plate zonal structures such as resting, proliferative, and hypertrophic chondrocytes as well as bone (**Figure 1A**) while the standard method mostly yielded bone tissues (**Figure 1B**). On the other hand, physeal drill debris generated during both procedures and also collected for analysis consisted of a heterogeneous mixture of cells and tissue fragments (**Figure 1C**).

**Figure 1.**
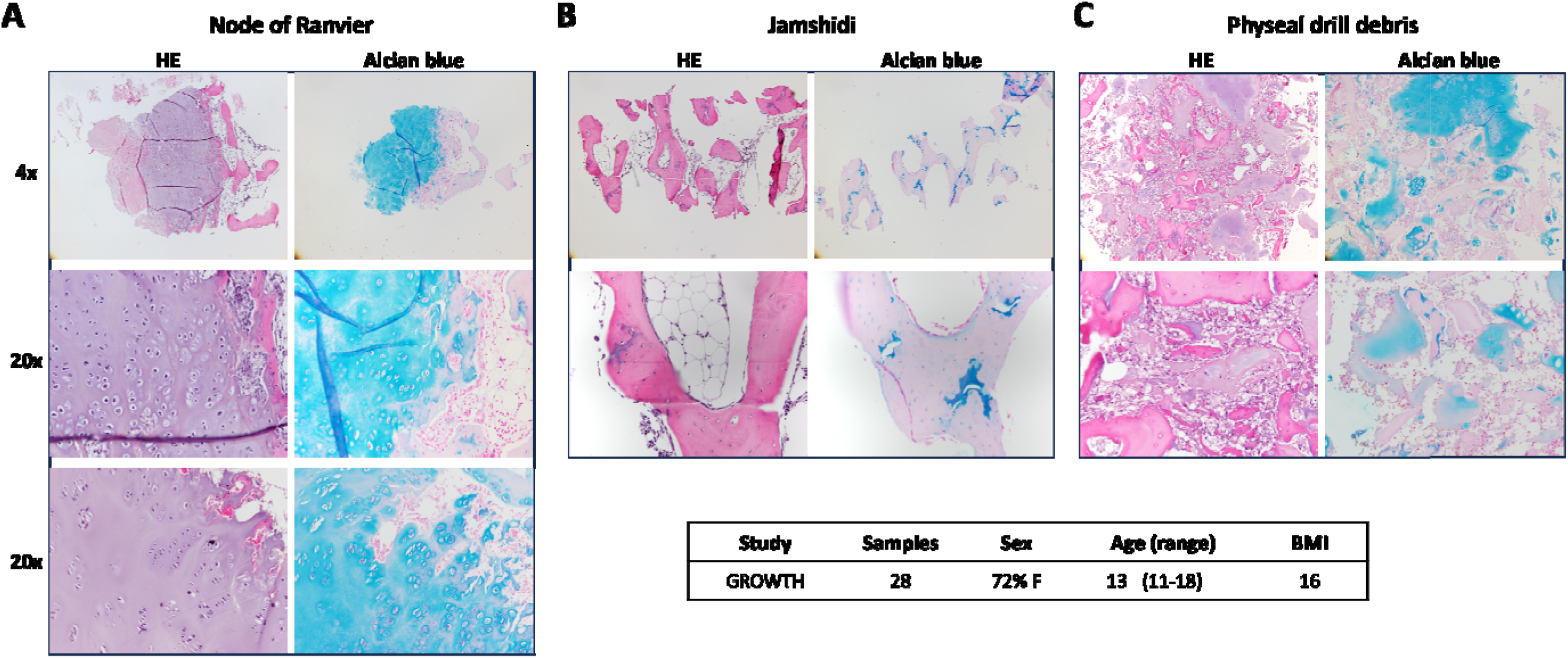
H&E and Alcian Blue staining of collected tissues using different biopsy methods. (**A**) Leiden method, clearly highlighting growth plate cartilage and bone structures. Note the characteristic columnar organization of chondrocytes in the lower panel. (**B**) Standard PE method, yielding mostly bone structures. (**C**) Physeal drill debris, displaying a heterogenous mixture of cells and tissue fragments.

### Growth plate chondrocyte characterization

Next, we compared *in vitro* cellular characteristics of growth plate and articular cartilage chondrocytes. The latter isolated from macroscopically preserved cartilage from our RAAK study (*25, 34*). To assess 2D proliferation, growth plate and articular chondrocytes were seeded and cell counts were performed after 24 and 96 hours of incubation. As shown in **Figure 2**, after 24 hrs the number of growth plate chondrocytes was five times higher than the number of articular chondrocytes. However, at 96 hours the number of cells appeared almost equal. The plot suggests both a cell cycle arrest of growth plate chondrocytes and a catch up growth of articular chondrocyte at a plateau of around 50x10^4^ cells, between 24 hrs and 96 hrs.

**Figure 2.**
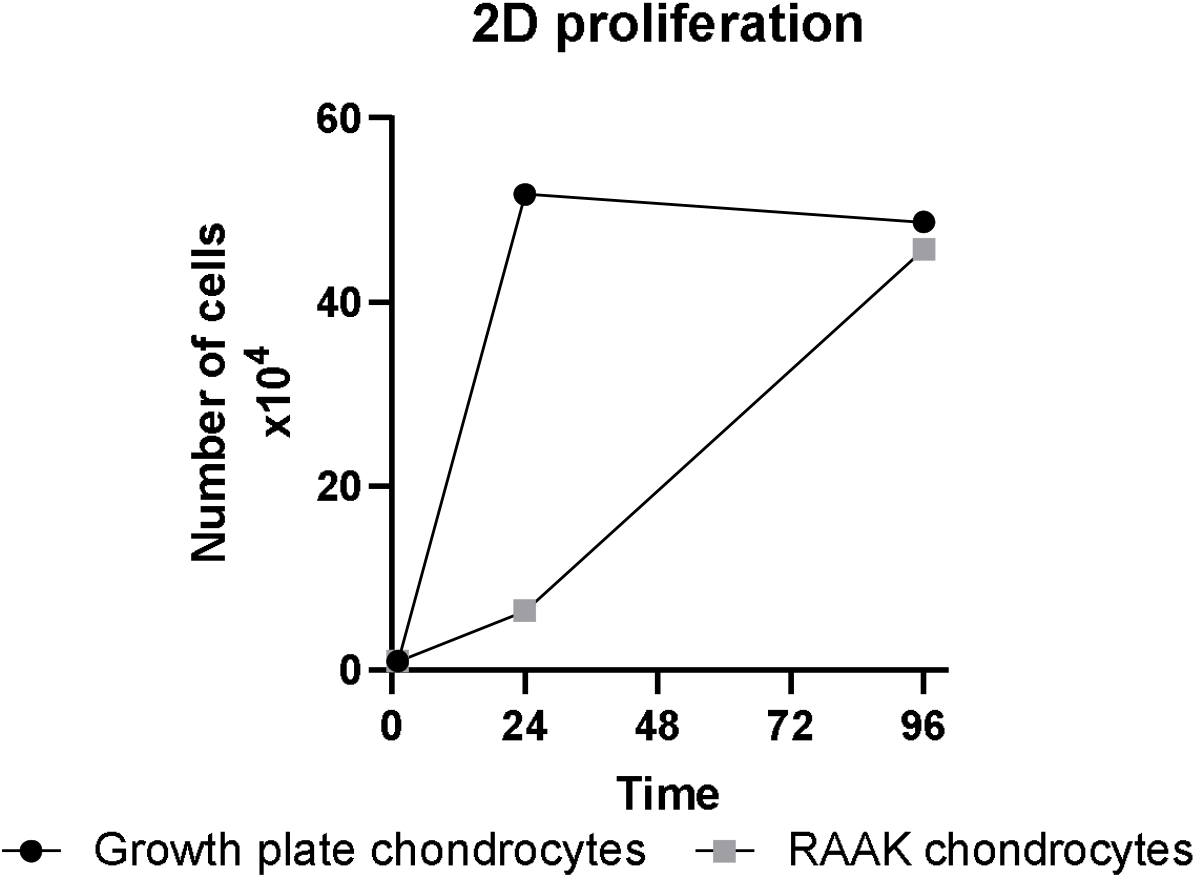
2D proliferation rate. Growth plate chondrocytes compared to OA chondrocytes (N=1).

Next, we generated 3D neo-cartilage organoids *in vitro*, from growth plate and OA chondrocytes to evaluate the ability to produce cartilage matrix. As illustrated in **Figure 3A**, growth plate chondrocytes formed welldefined spherical 3D cartilage organoids that were on average ∼2.1 mm in diameter whereas articular cartilage cells were generally smaller on average ∼1.5 mm in diameter. The cartilage matrix produced by growth plate chondrocytes exhibited Alcian Blue staining comparable to that of RAAK chondrocytes (**Figure 3B**). Finally, we measured the gene expression levels of cartilage markers *SOX9* and *COL2A1*, early hypertrophy marker *COL10A1*, and terminal maturation marker *MMP13*. As shown in **Figure 3C**, the growth plate derived organoids exhibited statistically significant higher expression of these genes compared to the articular cartilage derived organoids.

**Figure 3.**
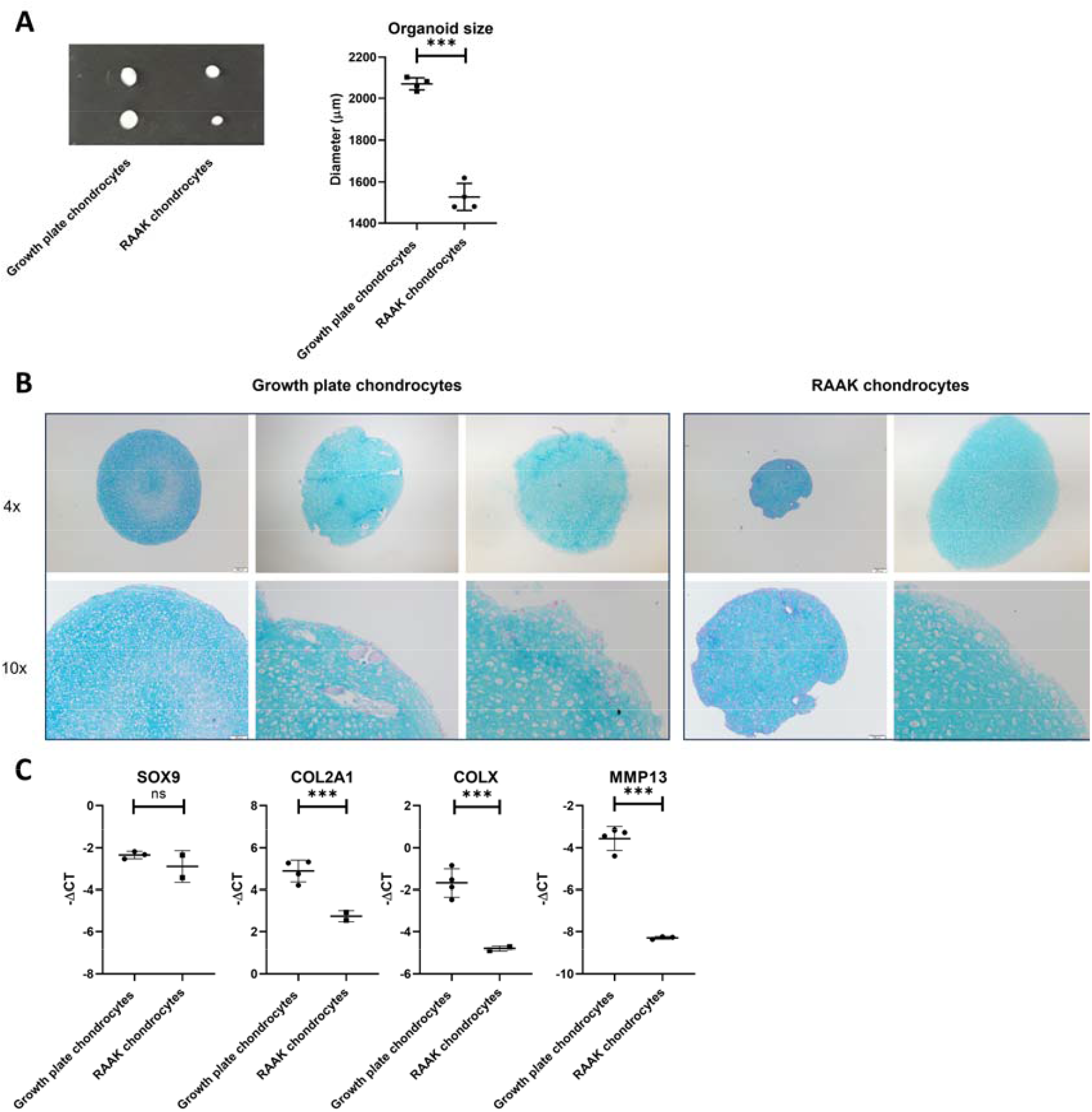
3D cartilage organoid characterization. (**A**) Size of 3D cartilage organoids. (**B**) Alcian Blue staining of 3D cartilage organoids. (**C**) Gene expression levels of 3D cartilage organoids. All measurements are done after 21 days of culturing. (N=1 donor, N=2-4 biological replicates, for both growth plate and RAAK chondrocytes).

## DISCUSSION

In this study, we demonstrate the feasibility and reliability of waste human growth plate tissues harvested from PE procedure for their use in functional, molecular epidemiological studies. Across multiple consecutive surgical procedures, our optimized surgical PE procedure confirmed consistently viable collection and storage of growth plate tissues and cells. Herein, histology of paraffin embedded samples confirmed that this growth plate tissue remained intact and with recognizable zonal growth plate chondrocyte states. Moreover, owing to established protocols (*24*), we were able to successfully isolate living growth plate chondrocytes. These deposited cartilage extracellular matrix resulting in 3D growth plate organoids thereby confirming the capacity of these cells to generate cartilage-like tissue.

The value of human growth plate collection is particularly clear when considering the current limitations in the field. Much of our understanding of human growth disorders is still based on germline genetic variants identified using DNA from peripheral blood leukocytes. While this approach has been highly informative to identify which genes were involved, it does not provide biological understanding of the mechanism at play in the growth plate. In the absence of human growth-plate tissues or cells, functional follow-up studies have been performed in mouse growth-plate chondrocytes, and provided insights (*14, 15, 35*). However, since there are important differences in timing and closure of rodent growth plates in comparison to that of humans (*17*), the data is poorly translated to humans hence drug target discoveries has been limited (*16*). We therefore advocate that the Leiden GROWTH study is bound to connect population-level genetic findings to the biological pathways that are active within the human growth plate. Moreover, application of our model allows to assess the potential impact of formally known but also of new variants of uncertain significance (VUS) such as those previously identified in *ACAN* (*36*). Noteworthy is that in comparing chondrocytes isolated from growth plate and articular cartilage, we found that growth plate chondrocytes had a significant larger initial proliferation and matrix deposition capacity. On the other hand, we showed that they vividly expressed markers of early hypertrophy (*CO10A1)* and terminal maturation (*MMP13*). These differences are likely attributed to an intrinsic difference in tissue origin and cell fate.

Potential limitation of our collection is that the PE referrals included growth plate from adolescents with underlying monogenetic syndromal etiology of tall stature, such as Marfan-syndrome. Nonetheless, since the LUMC is a nationally accredited expert centre for growth genetics and tall stature, inclusion criteria were, for each patient, unambiguously evaluated by expert specialists regarding signs of possible underlying monogenetic etiology of tall stature. This evaluation includes a thorough history, complete physical examination with anthropometric measurements (arm span, sitting height index, head circumference) and description of specific dysmorphic features. The remaining, over 90% of adolescents who are referred for a percutaneous epiphysiodesis have thus a common polygenic/non-syndromic form of tall stature, yet appear on the tall-statured end of the spectrum. Notably, herein is that identification of the relevant biological processes that determine chondrocyte transitions along the zonal growth plate structures, are captured by within sample maturation-states, hence do not suffer from a bias introduced when heterogeneous samples are compared. Moreover, adolescents with normal height undergoing surgery due to significant limb-length discrepancy can function as normal height control. This because these limb-length discrepancies occur to unilateral growth plate damage as a result of trauma or infection in the past. Henceforth, we do not expect that our study is subject to an inclusion bias of syndromic nor tall-stature only patients that could hamper future studies of physiologically relevant aspects of growth plate physiology.

Beyond better understanding of basic growth biology, the Leiden GROWTH study may also provide a valuable source for establishing human 3D organoid and/or explant tissue models of human growth plates, enabling reliable preclinical testing of compounds that target growth plate dysfunction, similar as outlined previously for articular cartilage (*26, 37*). Notably in this respect is Vosoritide, that has recently been identified and successfully targets aberrant function of the height gene *FGFR3* in short stature children with Achondroplasia (*38-40*). We anticipate that Vosoritide can function in these models as an established modifier of growth plate chondrocyte states hence a reliable validation of readout. Taken together, after decades of GH-dominated research and therapy for growth in limited patient groups the established and optimized ‘Leiden GROWTH study’ is bound to contribute to the advancement of personalized medicine in pediatric growth disorders.

## ACKNOWLEDGEMENT

We thank all study participants of the Leiden GROEI-study. Moreover, we thank everybody from the group who contributed to the collection of the joint tissues.

The Leiden University Medical Center has and is supporting the Leiden GROEI study. Furthermore, the research leading to the Leiden GROEI collection has received funding from the Netherlands Orthopedic Research and Education Funds (Anna Fonds, Grant number OZ.2019.22).

## Supplementary Material

**Supplementary Figure S1.**
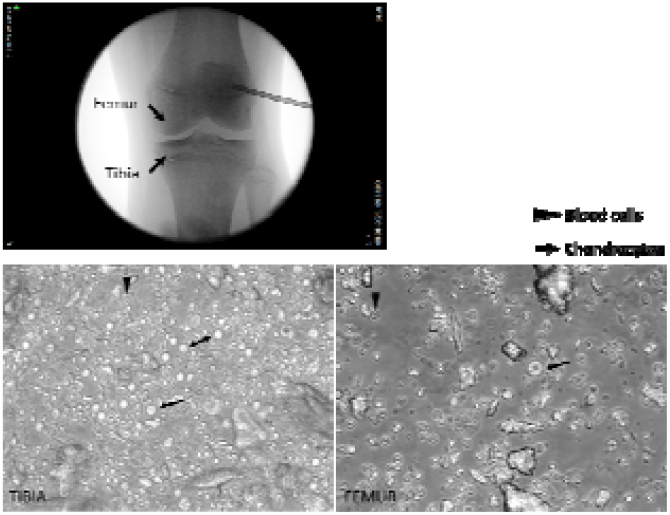
Surgical ablation of physis to stop growth; percutaneous epiphysiodesis (PE). Upper panel: needle angle for biopsy of growth plate with Jamshidi needle and entrance site for drill. Lower panel: representative light microscopy image of physeal drill debris collected from tibia and femur.

**Supplementary Table S1.**
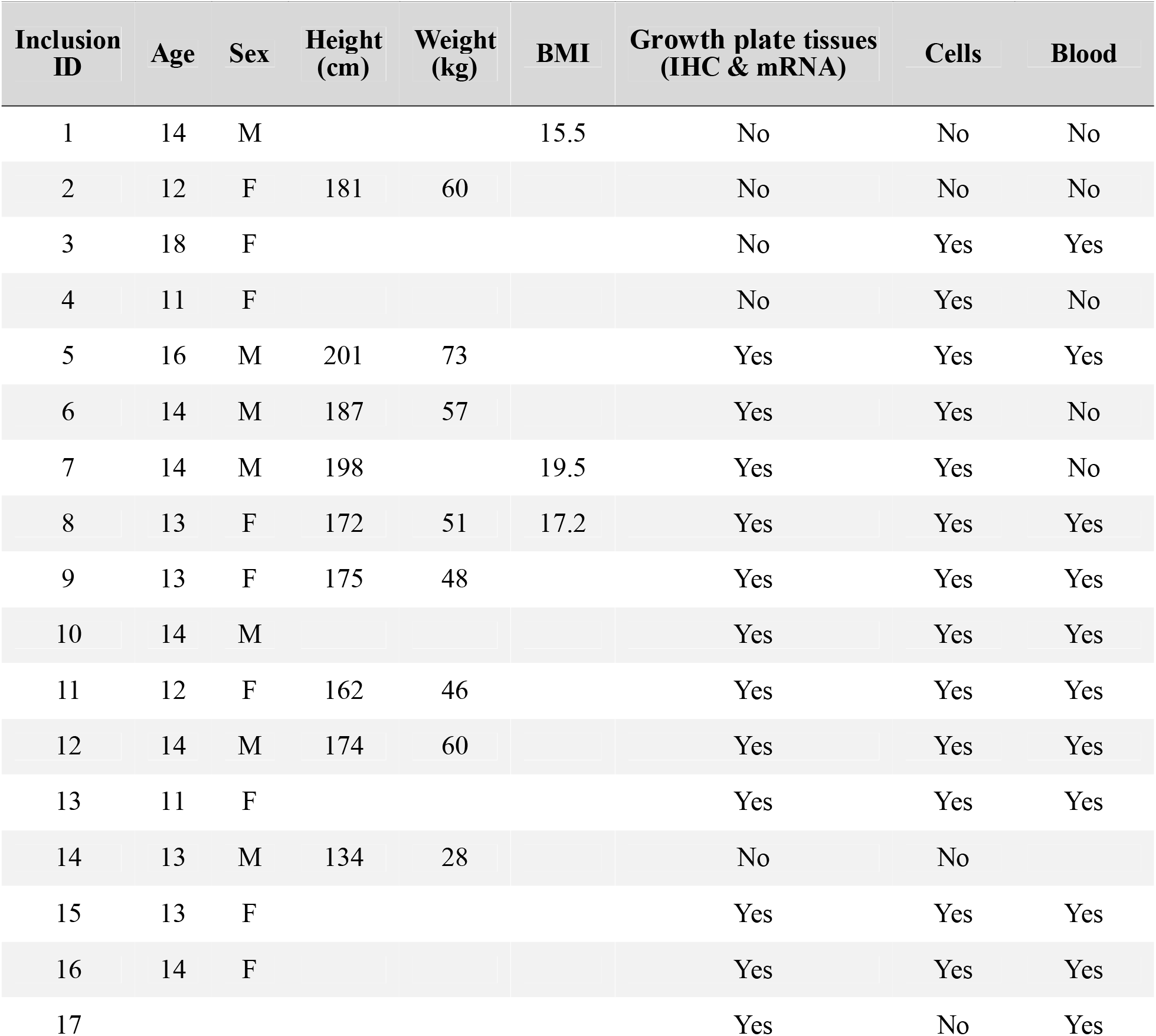
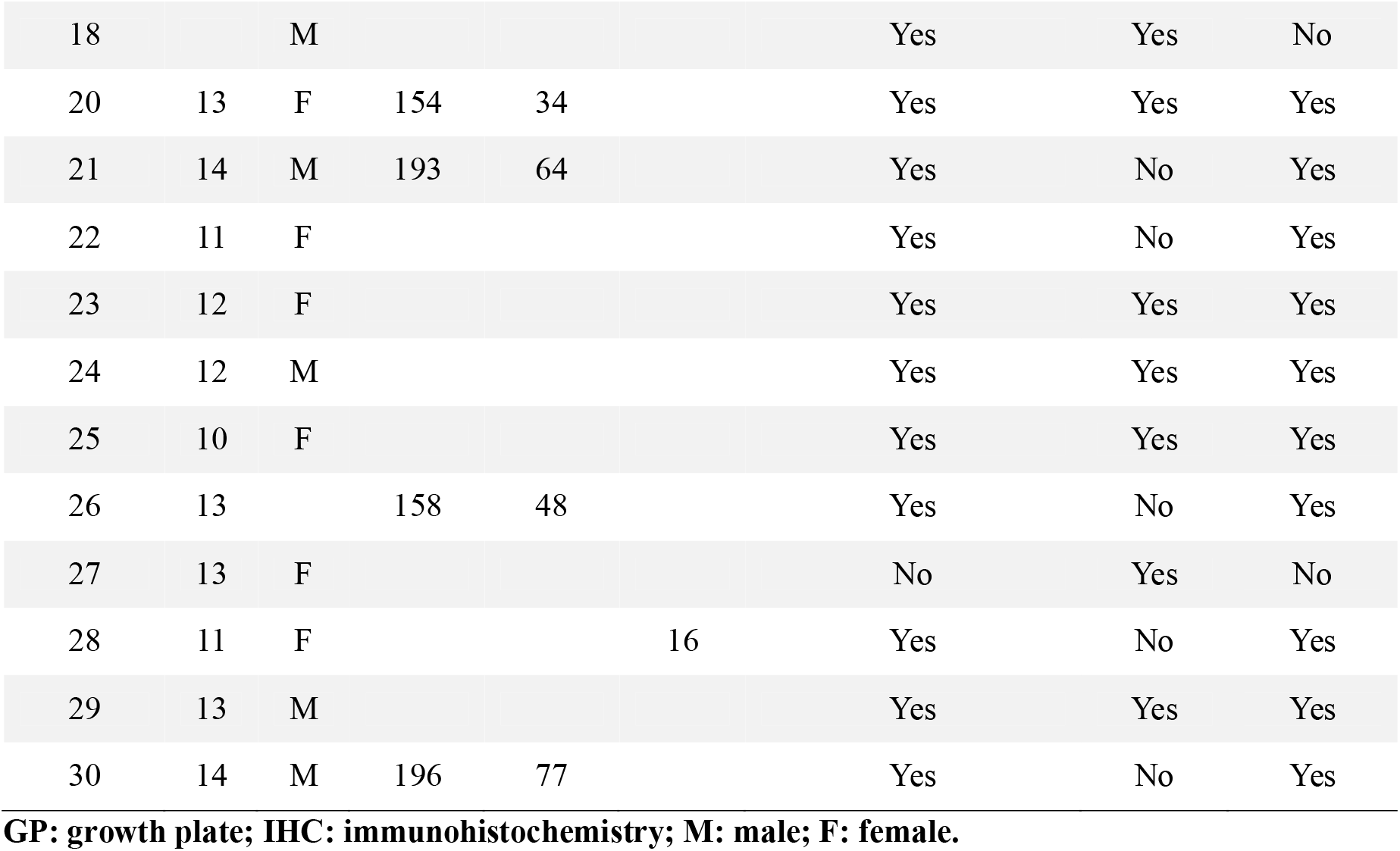
Patient characteristics and collection of biospecimen.

## REFERENCES

1. P. F. Collett-Solberg et al., Growth hormone therapy in children; research and practice–a review. Growth Horm. IGF Res. 44, 20–32 (2019).

2. R. Rapaport, J. M. Wit, M. O. Savage, Growth failure:’idiopathic’only after a detailed diagnostic evaluation. Endocrine Connections 10, R125–R138 (2021).

3. K. C. Song et al., Etiologies and characteristics of children with chief complaint of short stature. Annals of Pediatric Endocrinology & Metabolism 20, 34–39 (2015).

4. E. Bellotto et al., Pattern and features of pediatric endocrinology referrals: a retrospective study in a single tertiary center in Italy. Frontiers in Pediatrics 8, 580588 (2020).

5. B. Villafuerte, R. Barrio, M. Martín-Frías, M. Alonso, B. Roldán, Auxological characteristics of pediatric patients with permanent or transient isolated growth hormone deficiency. Response to treatment and final height. Endocrinología, Diabetes y Nutrición (English ed.) 66, 368–375 (2019).

6. A.-B. Ariza-Jimenez, I. Leiva Gea, M. J. Martinez-Aedo Ollero, J. P. Lopez-Siguero, Isolated growth hormone deficiency and idiopathic short stature: comparative efficiency after growth hormone treatment up to adult height. Journal of Clinical Medicine 10, 4988 (2021).

7. A. D. Rogol, G. F. Hayden, Etiologies and early diagnosis of short stature and growth failure in children and adolescents. The Journal of pediatrics 164, S1-S14. e16 (2014).

8. N. L. M. Andrade et al., Diagnostic yield of a multigene sequencing approach in children classified as idiopathic short stature. Endocrine Connections 11, (2022).

9. E. V. Albuquerque, R. C. Scalco, A. A. Jorge, Management of endocrine disease: diagnostic and therapeutic approach of tall stature. European Journal of Endocrinology 176, R339–R353 (2017).

10. P. Wainschtein et al., Assessing the contribution of rare variants to complex trait heritability from whole-genome sequence data. Nat. Genet. 54, 263–273 (2022).

11. A. R. Wood et al., Defining the role of common variation in the genomic and biological architecture of adult human height. Nat Genet 46, 1173–1186 (2014).

12. L. Yengo et al., A saturated map of common genetic variants associated with human height. Nature 610, 704–712 (2022).

13. E. Cano-Gamez, G. Trynka, From GWAS to Function: Using Functional Genomics to Identify the Mechanisms Underlying Complex Diseases. Frontiers in Genetics 11, (2020).

14. M. Guo et al., Epigenetic profiling of growth plate chondrocytes sheds insight into regulatory genetic variation influencing height. eLife 6, e29329 (2017).

15. M. Wuelling et al., Epigenetic Mechanisms Mediating Cell State Transitions in Chondrocytes. J. Bone Miner. Res. 36, 968–985 (2021).

16. J. Baron et al., Short and tall stature: a new paradigm emerges. Nature Reviews Endocrinology 11, 735–746 (2015).

17. S. H. Kilborn, G. Trudel, H. Uhthoff, Review of growth plate closure compared with age at sexual maturity and lifespan in laboratory animals. Contemp Top Lab Anim Sci 41, 21–26 (2002).

18. E. Hunziker, R. Schenk, L. Cruz-Orive, Quantitation of chondrocyte performance in growth-plate cartilage during longitudinal bone growth. Jbjs 69, 162–173 (1987).

19. R. T. Ballock, R. J. O’Keefe, Physiology and pathophysiology of the growth plate. Birth Defects Research Part C: Embryo Today: Reviews 69, 123–143 (2003).

20. C. L. Peagler Jr, A. J. Dobek, S. Tabaie, S. A. Tabaie, Trends in the use of total hip arthroplasty in the pediatric population: a review of the literature. Cureus 15, (2023).

21. M. Jäger et al., Primary total hip replacement in childhood, adolescence and young patients: quality and outcome of clinical studies. Technol. Health Care 16, 195–214 (2008).

22. E. Benyi et al., Efficacy and safety of percutaneous epiphysiodesis operation around the knee to reduce adult height in extremely tall adolescent girls and boys. Int. J. Pediatr. Endocrinol. 2010, 740629 (2010).

23. W. J. Goedegebuure et al., Long-term follow-up after bilateral percutaneous epiphysiodesis around the knee to reduce excessive predicted final height. Arch. Dis. Child. 103, 219–223 (2018).

24. Y. F. Ramos et al., Genes involved in the osteoarthritis process identified through genome wide expression analysis in articular cartilage; the RAAK study. PLoS One 9, e103056 (2014).

25. M. Tuerlings et al., WWP2 confers risk to osteoarthritis by affecting cartilage matrix deposition via hypoxia associated genes. Osteoarthritis Cartilage 31, 39–48 (2023).

26. E. Houtman et al., Human Osteochondral Explants: Reliable Biomimetic Models to Investigate Disease Mechanisms and Develop Personalized Treatments for Osteoarthritis. Rheumatol Ther 8, 499–515 (2021).

27. R. Coutinho de Almeida et al., Identification and characterization of two consistent osteoarthritis subtypes by transcriptome and clinical data integration. Rheumatology 60, 1166–1175 (2021).

28. Y. Schönbeck et al., Increase in prevalence of overweight in Dutch children and adolescents: a comparison of nationwide growth studies in 1980, 1997 and 2009. PLoS One 6, e27608 (2011).

29. A. M. Fredriks, S. Van Buuren, W. Van Heel, R. Dijkman-Neerincx Hm; Verloove-Vanhorick, S. & Wit, JM Nationwide age references for sitting height, leg length, and sitting height/height ratio, and their diagnostic value for disproportionate growth disorders. Arch. Dis. Child 90, 807–812 (2005).

30. P. van Dommelen, Y. Schönbeck, S. van Buuren, A simple calculation of the target height. Arch. Dis. Child. 97, 182–182 (2012).

31. H. H. Thodberg, O. G. Jenni, J. Caflisch, M. B. Ranke, D. D. Martin, Prediction of adult height based on automated determination of bone age. The Journal of Clinical Endocrinology & Metabolism 94, 4868–4874 (2009).

32. P. S. Greulich WW, Radiographic Atlas of Skeletal Development of the Hand and Wrist. . n. edition., Ed., (Stanford University Press, CA:, 1959).

33. N. Bomer et al., Aberrant Calreticulin Expression in Articular Cartilage of Dio2 Deficient Mice. PLoS One 11, e0154999 (2016).

34. R. G. M. Timmermans et al., A human in vitro 3D neo-cartilage model to explore the response of OA risk genes to hyper-physiological mechanical stress. Osteoarthr Cartil Open 4, 100231 (2022).

35. T. D. Capellini et al., Ancient selection for derived alleles at a GDF5 enhancer influencing human growth and osteoarthritis risk. Nat Genet 49, 1202–1210 (2017).

36. J. S. Renes et al., Clinical characteristics of pathogenic ACAN variants and 3-year response to growth hormone treatment: real-world data. Horm. Res. Paediatr. 97, 456–469 (2024).

37. E. Houtman et al., Inhibiting thyroid activation in aged human explants prevents mechanical induced detrimental signalling by mitigating metabolic processes. Rheumatology 62, 457–466 (2022).

38. R. Savarirayan et al., Phase 3 Trial of Oral Infigratinib in Children with Achondroplasia. N. Engl. J. Med., (2026).

39. O. Semler, H. Hoyer-Kuhn, Navepegritide in children with achondroplasia. Translational Pediatrics 15, 260 (2026).

40. M. A. Kalapurackal, P. Wanker, Early initiation of therapy with vosoritide: case report. Description of the first Italian patient affected by achondroplasia treated with vosoritide before 2 years of age. Frontiers in Pediatrics 14, 1741018 (2026).

